# Compost-associated microbiota modulates root transcriptional responses and enhances tomato drought resilience

**DOI:** 10.64898/2026.10.29.762138

**Authors:** Rosalba Cipriani, Susan R. Strickler, Francesco Petruzzellis, Giulia Ghirardello, Leonardo Greggio, Werner Zanardi, Tiziano Bonato, Barbara Baldan, Sebastiano Nigris

## Abstract

- Compost is a promising sustainable alternative to mineral fertilization because it supplies nutrients and organic matter while hosting diverse microbial communities. However, even if its application to plants has been proven to alleviate different biotic and abiotic stresses, it remains unclear whether its benefits arise from its physico-chemical properties, its associated microbiota, or both.
- In this study, we compared the impact of compost, sterile compost and an NPK treatment providing equivalent macronutrient inputs on tomato plants cv. Micro-Tom. After assessing physiological parameters of the leaves and molecular responses of the roots under optimal conditions, we performed a drought stress experiment under the three fertilization treatments, complementing the analysis with the investigation of substrate water-retention properties.
- Although all fertilized treatments improved plant performance relative to nutrient-poor soil, RNA sequencing revealed distinct root transcriptional profiles among compost, sterile compost and NPK, with the compost treatment upregulating several pathways related to plant growth and health, such as hormone signaling, nutrient metabolism, and stress responses.
- When subjected to severe water stress, compost-treated plants retained the highest photosynthetic efficiency and compost showed the highest available water content compared to the other treatments. These findings suggest that compost may enhance tomato drought resilience through combined effects of substrate water availability and compost-associated microbiota.

## 1. Introduction

Drought is a global concern and one of the major threats to agriculture, as crop production depends on water, soil, and climate and is therefore highly susceptible to the increasing frequency and intensity of water stress. Between 1991 and 2023, droughts caused an estimated USD 278 billion in agricultural losses worldwide. However, the actual losses were probably higher because drought impacts are often underreported (FAO Report, 2025). Agriculture accounts for approximately 82% of the total impact of drought, compared to 18% across all other sectors, highlighting its particular vulnerability (FAO Report, 2021; Li *et al*., 2026). At the same time, the global population is expected to increase by approximately 24% over the next 50 years (UN population prospects, 2024), while global food demand is projected to increase by 35% to 56% between 2010 and 2050 (van Dijk *et al*., 2021). Meeting this growing demand therefore requires sustained improvements in crop productivity. This need has contributed to the widespread use of chemical fertilizers, which provide readily available nutrients but can negatively affect soil health when used intensively over long periods, promoting compaction and acidification while reducing soil organic matter, microbial diversity, and microbial enzyme activity (Pahalvi *et al*., 2021). These effects highlight the need for alternative fertilization strategies that sustain plant growth while preserving soil health and improving resilience to water limitation, particularly under severe drought conditions.

In recent decades, organic amendments have therefore attracted increasing attention as more sustainable alternatives, due to their positive effects on soil structure, fertility, and microbial communities (Matisic *et al*., 2024). Beyond these benefits, organic fertilizers can also enhance plant tolerance to abiotic stress. By improving soil water retention and nutrient availability, they can help maintain plant water status, photosynthetic activity, membrane stability, and growth during drought (Liu *et al*., 2024; Mehmood *et al*., 2025). Organic amendments may also modulate osmotic adjustment and antioxidant responses, thereby limiting oxidative damage under stress. Their beneficial effects have also been reported in saline and metal-contaminated soils, where they can improve ion balance, reduce the bioavailability of toxic elements, and support the activity of beneficial soil microorganisms (Irin and Hasanuzzaman, 2024). However, the magnitude and mechanism of these effects depend on the amendment type, soil properties, crop species, and stress intensity.

Among organic amendments, compost is one of the oldest and most widely used fertilizers. Compost provides organic matter and nutrients and hosts a diverse microbial community, mainly composed of bacteria and fungi, which can contribute to soil fertility and plant health. Several taxa commonly found in compost, including members of the Firmicutes, Actinobacteria, Proteobacteria, Bacteroidetes, and Chloroflexi, have been associated with plant growth promotion, enhanced nutrient acquisition, phosphate solubilization, nitrogen fixation, and suppression of phytopathogens (Aguilar-Paredes *et al*., 2023). Following application to soil, compost can also reshape microbial community structure and interaction networks, contributing to soil and plant performance under both optimal and stressful conditions (Heisey *et al*., 2022). However, despite the recognized benefits of compost, the relative contribution of its living microbial component compared with its organic matter and nutrient content remains poorly understood. In particular, it remains unclear to what extent the impact of compost on plant physiological and molecular responses result from its physicochemical properties and nutrient supply, from the activity of its associated microbiota, or from their combined effect.

Among the crops particularly affected by drought is tomato (*Solanum lycopersicum* L.), an economically important horticultural crop. Water limitation can impair plant water status and photosynthesis, as well as flower development and pollen fertility, ultimately reducing fruit set, yield, and quality (Al Achkar *et al*., 2026). Furthermore, cultivated tomato species have relatively limited genetic diversity for drought-related traits, which are instead more extensively present in their wild relatives (Krishna *et al*., 2019). There is therefore a need for sustainable strategies capable of supporting tomato productivity and health under water-limited conditions. Tomato is also a suitable model for investigating plant responses to environmental and nutritional conditions because of its fully sequenced genome, extensive genetic and genomic resources, broad cultivar diversity, and the availability of short-cycle genotypes such as Micro-Tom (Conti *et al*., 2023).

In this study, we investigated the contribution of compost-associated microbiota to plant responses under both physiological and drought stress conditions using an industrially produced compost derived from organic food waste (biowaste). To distinguish, as far as possible, the effects of the living microbial component from those associated with compost-derived organic matter and nutrient supply, we compared three treatments: compost, sterilized compost, and a home-made NPK fertilizer providing amounts of macronutrients equivalent to those provided by compost. This experimental design allowed us to assess plant physiological and molecular responses under optimal conditions and to determine (i) how these responses changed under increasing drought severity, and (ii) whether the presence of compost-associated microbiota contributes to plant performance beyond the nutritional and physico-chemical effects of compost.

## 2. Materials and methods

### 2.1 Soil treatments

The baseline soil used in the experiments consisted of 45% agricultural field soil, 45% peat, 5% perlite, and 5% sand. Its physico-chemical properties were determined according to UNI standard methods (Table S1). The compost used in this work was produced by S.E.S.A. S.p.A. (Este, Italy), and its physico-chemical properties were determined according to UNI standard methods and were in line with EU directives and consistent with the company standards (Table S2).

Sterile compost was obtained by autoclaving at 121 °C for 2 h. Sterility was assessed by suspending 10 g of Sterile Compost in 50 mL of sterile deionized water and shaking the suspension on an orbital shaker for 1 h. An aliquot of 100 µL was spread onto Plate Count Agar (PCA) and incubated at 28 °C for 4 days. No colonies were detected under these culture conditions, confirming successful sterilization. To verify any possible effect of the sterilization process on plant substrate, the pH of soil/compost and soil/sterile compost mixtures was measured, with no significant difference observed.

The three soil treatments were prepared by mixing the baseline soil with either 33 g/L compost, 33 g/L sterile compost, or the corresponding amounts of NPK fertilizers. For the NPK treatment, nitrogen, phosphorus, and potassium were supplied using urea, diammonium phosphate (DAP), and potassium chloride (KCl), respectively, to match the macronutrient amounts provided by the compost. The final application rates were 0.794 g/kg DAP, 0.311 g/kg urea, and 0.882 g/kg KCl. All amendments were accurately weighed, thoroughly mixed with the soil, and subsequently allocated into 0.5-L pots.

### 2.2 Plant growth

Tomato seeds (*Solanum lycopersicum* L.) cv. Micro-Tom were surface-sterilized in a 5% sodium hypochlorite solution on an orbital shaker for 20 min, followed by four washes with sterile water. After 48 h of imbibition at room temperature, the seeds were sown on half-strength Murashige and Skoog (MS) basal medium supplemented with 1.5% agar and germinated at 25 °C and 65% relative humidity under a 16 h light/8 h dark photoperiod. Five days after germination, seedlings were transplanted into 0.5-L pots containing the respective soil treatments and returned to the growth chamber under the same environmental conditions. Pots were randomly arranged to minimize position-dependent effects. All plants were watered uniformly and regularly throughout the growth period.

### 2.3 Experimental design for growth assessment and transcriptomic analysis

For the growth test and RNA-sequencing experiment, 40 plants were initially established and equally distributed among the four substrate treatments: Soil, Compost, Sterile Compost, and NPK. Plants were grown in a growth chamber for one month, after which photosynthetic efficiency (F_v_’/F_m_’) was measured using a MultispeQ v2.0 device, and leaf dry biomass was determined. F_v_’/F_m_’ was measured approximately twice per plant, resulting in a total of 68 measurements.

For transcriptomic analysis, roots from five independent plants per substrate (Compost, NPK, and Sterile Compost) were collected, resulting in 15 biological samples for RNA sequencing. The experimental design is summarized in the figure below.

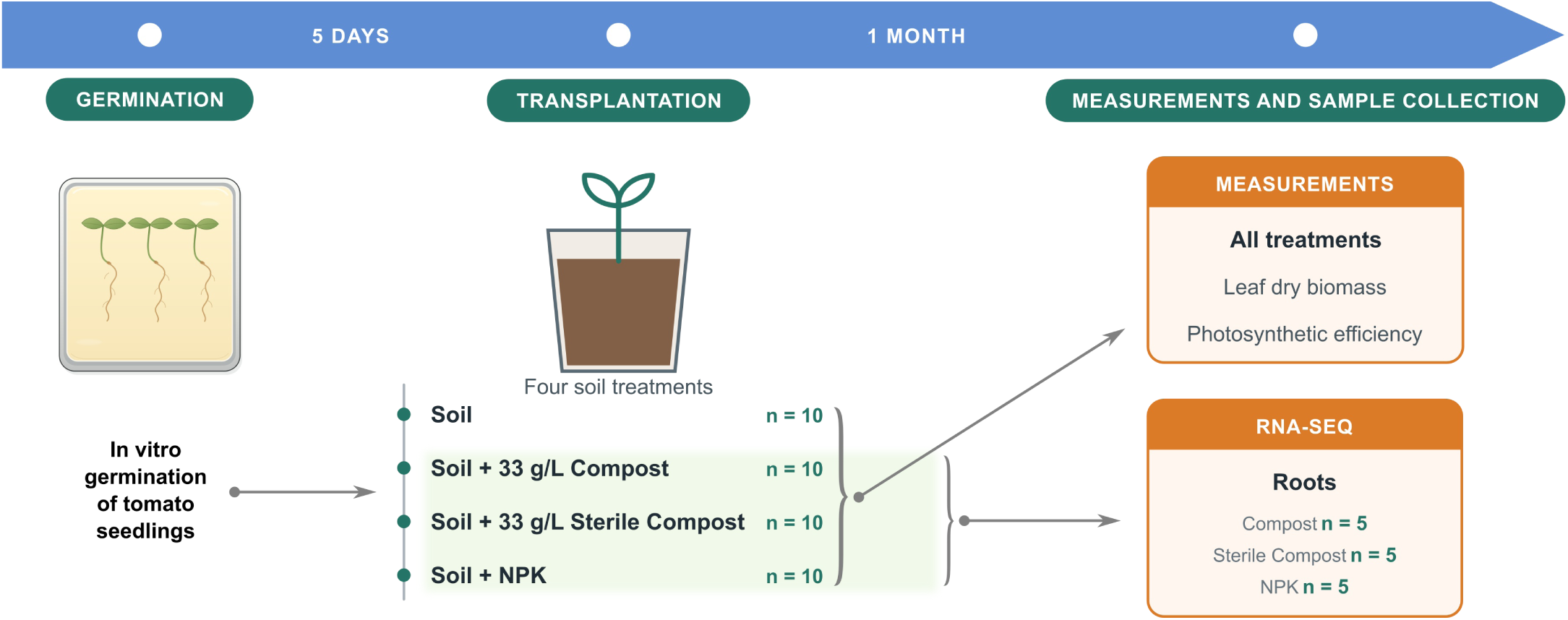

### 2.4 RNA extraction and transcriptome sequencing

After one month of growth, roots from 15 randomly selected tomato plants grown with Compost, Sterile Compost and NPK (5 samples per treatment) were collected, washed to remove adhering soil, and immediately frozen in liquid nitrogen. The tissues were homogenized using a mortar and pestle under liquid nitrogen, and total RNA was extracted using the CTAB method (Chang *et al*., 1993), with the modification of adding one-third volume of 8 M LiCl to the aqueous phase to obtain a final concentration of 2 M LiCl. RNA concentration and purity were assessed using an Implen NanoPhotometer N60 (Implen GmbH, Munich, Germany), while RNA integrity was evaluated by electrophoresis on a 1.2% agarose gel following RNA denaturation at 65 °C for 5 min.

RNA samples were sequenced by Novogene (UK) Co. The provider performed RNA quality control, poly(A)-enriched mRNA library preparation, sequencing on an Illumina NovaSeq X Plus platform (paired- end 150 bp; ∼12 Gb raw data per sample), and data quality control.

### 2.5 Transcriptomic analysis

Read quality was assessed using FastQC v0.12.1 and summarized across samples using MultiQC v1.27.1. Raw paired-end RNA-seq reads were processed using a custom Bash pipeline. First, ribosomal RNA reads were removed using SortMeRNA v4.3.7 (Kopylova *et al*., 2012), with the default SortMeRNA database. The remaining reads were quality-filtered and adapter-trimmed using Trimmomatic v0.39 (Bolger *et al*., 2014). Paired-end reads were processed using the following parameters: ILLUMINACLIP:TruSeq3-PE.fa:2:30:10, LEADING:5, TRAILING:5, SLIDINGWINDOW:4:15, and MINLEN:50. Quality-filtered reads were aligned to the *Solanum lycopersicum* reference genome SL4.0 using HISAT2 v2.2.1 (Kim *et al*., 2019), using the corresponding HISAT2 genome index. The resulting SAM files were converted to BAM format and sorted using SAMtools v1.13 (Li *et al*., 2009). Transcript-level expression was quantified from the sorted BAM files using StringTie v3.0 (Pertea *et al*., 2015). Quantification was performed against the *S. lycopersicum* ITAG4.1 gene annotation. Gene- and transcript-level count matrices were subsequently generated using the prepDE.py script distributed with StringTie, producing gene and transcript count matrices for downstream differential expression analysis. The resulting gene count matrix was imported into R and analyzed using the *edgeR* package v4.10.7 (Robinson et al., 2010).

### 2.6 Experimental design for drought stress

Tomato plants grown for three weeks under three fertilization conditions: Compost, Sterile Compost, and NPK were transferred to autoclaved peat pots to allow better control of soil moisture and were subsequently subjected to different levels of water availability for 14 days. A total of 300 plants were initially established for the experiment.

Water stress was performed by varying the volume of deionized water supplied to each plant every two days over the two-week experimental period. Deionized water was used to avoid introducing additional salts or nutrients that could interfere with the nutrients supplied by the different fertilization treatments. The water volumes used for the drought treatments were established in two preliminary trials conducted prior to the main experiment, in which different water amounts were evaluated to identify conditions producing distinct levels of water stress. Based on these preliminary trials, the following treatments for each pot were selected:

- Control: full hydration with deionized water;
- Low Stress: 35 mL of deionized water every two days;
- High Stress: 20 mL of deionized water every two days.

The experimental design is summarized in the figure below.

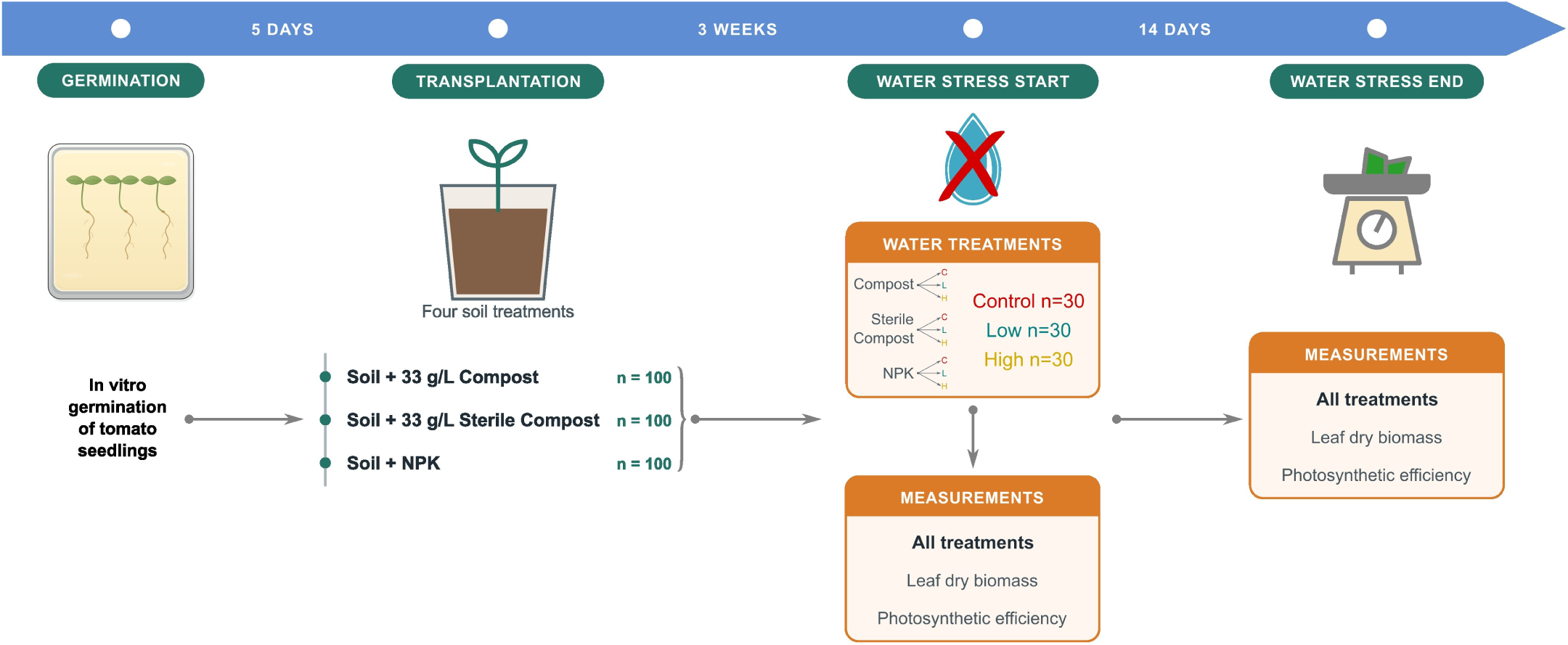

### 2.7 Measurements of physiological parameters under drought stress

After transferring plants in peat pots, before starting the water stress experiment (day 0) leaf dry biomass was measured for 131 plants (45 Compost, 45 Sterile Compost, and 41 NPK), while at day 14 it was measured for 146 plants (50 Compost, 52 Sterile Compost, and 44 NPK). The day-14 plants were distributed across Control, Low stress, and High stress water treatments as described above. Differences in sample size between time points reflect plant mortality during the experiment.

Photosynthetic performance was assessed by measuring the maximum photochemical efficiency of photosystem II in the light-adapted state (F_v_’/F_m_’) using a MultispeQ v2.0 device. Measurements were performed at the beginning of the experiment (day 0) and after 14 days of water stress. F_v_’/F_m_’ was measured on the same plants used for biomass determination at each time point. Approximately two or three F_v_’/F_m_’ measurements were obtained per plant at each sampling time. At day 0, 347 F_v_’/F_m_’ measurements were obtained from 131 plants, while at day 14, 320 measurements were obtained from 146 plants.

### 2.8 Statistical analysis of physiological parameters

For the initial growth test, differences among substrate treatments in F_v_’/F_m_’ and leaf biomass were evaluated separately using Kruskal-Wallis tests in R version 4.6.1 (R Core Team, 2026), followed by Dunn’s post hoc pairwise comparisons. P-values were adjusted for multiple comparisons using the Benjamini-Hochberg method (Benjamini and Hochberg, 1995), and statistical significance was set at α = 0.05.

For the drought stress experiment, F_v_’/F_m_’ and dry biomass were analyzed separately for each sampling day. Because F_v_’/F_m_’ is a continuous proportion bounded between 0 and 1, beta regression models with a logit link function were fitted using the *betareg* R package v3.2-6 (Cribari-Neto and Zeileis, 2010). The mean component included fertilization, water stress, and their interaction, while the dispersion component was modelled as a function of the fertilization × water stress interaction. The model including the interaction in both components was selected based on the lowest Akaike Information Criterion (AIC; Akaike, 1974). Dry biomass was analysed using Gamma regression models with a log link, fitted with the *glmmTMB* package v1.1.15.2 (Brooks *et al*., 2017). The models included fertilization, water stress, and their interaction in the mean component. Candidate models with and without group-specific dispersion were compared using AIC. At day 0, the final biomass model included the fertilization × water stress interaction in the mean component but not in the dispersion component, whereas at day 14 the interaction was included in both components. For both response variables, model adequacy was assessed using residual diagnostics implemented in the *DHARMa* package v0.5.0 (Hartig *et al*., 2026). Estimated marginal means were calculated on the response scale using the *emmeans* package v2.0.4 (Lenth, 2026), and pairwise comparisons were performed among water stress levels within each fertilization treatment and among fertilization treatments within each water stress level. P-values were adjusted using the Benjamini-Hochberg method, with statistical significance set at α = 0.05.

### 2.9 Soil available water content assessment

#### 2.9.1 Soil moisture release curves

Soil moisture release curves were measured in order to derive the Available Water Content (AWC) of each soil treatment (i.e., Soil, Soil + 33 g/L of compost, Soil + 33 g/L of sterile compost, Soil + NPK). The procedure was adapted from Nardini *et al*. (2021).

For each substrate, approximately 100 mL of substrate was brought to field capacity by gradually adding tap water while mixing until no further drainage was observed. Six samples for every substrate of approximately 7 mL each were then prepared in pre-weighed disposable plastic cups (maximum capacity 15 mL) and weighed to determine their initial fresh weight (FW_0_). Samples were partially dehydrated to approximately 10% fresh mass loss to discard oversaturation water. The target mass was calculated as:

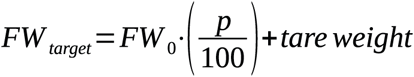

where FW_0_ is the initial fresh weight and p is the percentage of mass loss. Once the target mass was reached, samples were sealed and equilibrated for 30 minutes before measurement. After preparation, sequential measurements of soil water potential (Ψ) and corresponding fresh weight (FW) were taken during controlled dehydration. After each dehydration step, samples were sealed in the sample holder for 1 h before measuring soil Ψ. Soil Ψ was measured using a dew point potentiometer (WP4C, Meter Inc., Pullman, WA, USA), which determines Ψ by measuring the vapor pressure of the headspace in a closed chamber, where the liquid-phase of the sample and the vapor-phase in the chamber reach the equilibrium.

This procedure was repeated over five days, yielding six to ten measurements per aliquot and WP values ranging from ‒0.1 to ‒5.0 MPa. At the end of the measurements, the aliquots were oven-dried for 48 hours at 45 °C and subsequently weighed to obtain the dry weight (DW). Overall, three weeks were required to collect all the data for each soil substrate.

Soil water retention curves were obtained by relating soil Ψ to the corresponding gravimetric Water Content (WC), calculated as:

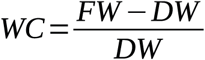

Experimental data were modeled using nonlinear regression to describe the behavior of the soil water retention curves. Two models (expoDecay and powerCurve) were compared using R software; the powerCurve model was selected based on lower Akaike Information Criterion (AIC) values.

#### 2.9.2 Available Water Content (AWC)

Available Water Content (AWC) was calculated as the difference between the saturated water content (SWC) and the water content at the Permanent Wilting Point (PWP; ‒1.5 MPa):

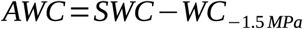

SWC was determined from three additional samples for each soil treatment, which were suspended in sealed vials, with the base filled with a 0.05 m KCl solution (Ψ = ‒0.23 MPa) for 24 h. Following this equilibration time, soil Ψ and FW were measured, as well as DW, after oven-drying samples for 48 h. WC at soil Ψ equal to ‒1.5 MPa was obtained from the corresponding moisture release curves.

Differences between AWC values among substrates were calculated as follows, using R software. A bootstrap procedure was applied to estimate mean values and the 2.5% and 97.5% confidence intervals (CIs) of AWC for each substrate. Specifically, a resampling procedure was performed by randomly sampling, with replacement, the same number of points as in the original dataset (corresponding to 100% of the available points for each substrate) from the moisture release curves. The selected subsets of points were then modelled as described above, and the corresponding AWC values were calculated for each substrate. This procedure was repeated 999 times, generating a distribution of AWC values for each substrate. Mean values and 2.5% and 97.5% CIs of AWC were subsequently derived from these distributions. Differences between substrates were considered significant when the CIs did not overlap.

## 3. Results

### 3.1 Tomato plant physiological performance is primarily determined by nutrient availability

To assess whether compost-associated microbiota affects tomato plant performance independently of nutritional inputs, plants were grown under four soil treatments for one month, after which leaf biomass and photosynthetic efficiency were measured (Figure 1): compost containing its native microbiota (Fig. 1B), sterilized compost (Fig. 1C), and chemical fertilization with NPK (Fig. 1D). Nutrient-poor soil was included as a baseline condition (Fig. 1A). After one month of growth, plants grown in all three fertilized conditions showed significantly greater leaf dry biomass than plants grown in nutrient-poor soil (Fig. 1E). However, the distribution of biomass differed among the fertilization treatments, with compost-grown plants showing a lower biomass than NPK-treated plants, while sterilized compost resulted in an intermediate phenotype. Photosynthetic efficiency (F_v_’/F_m_’) was also strongly reduced in nutrient-poor soil, whereas plants grown with compost, sterilized compost, or NPK maintained significantly higher and comparable photosynthetic efficiency (Fig. 1F).

**Figure 1.**
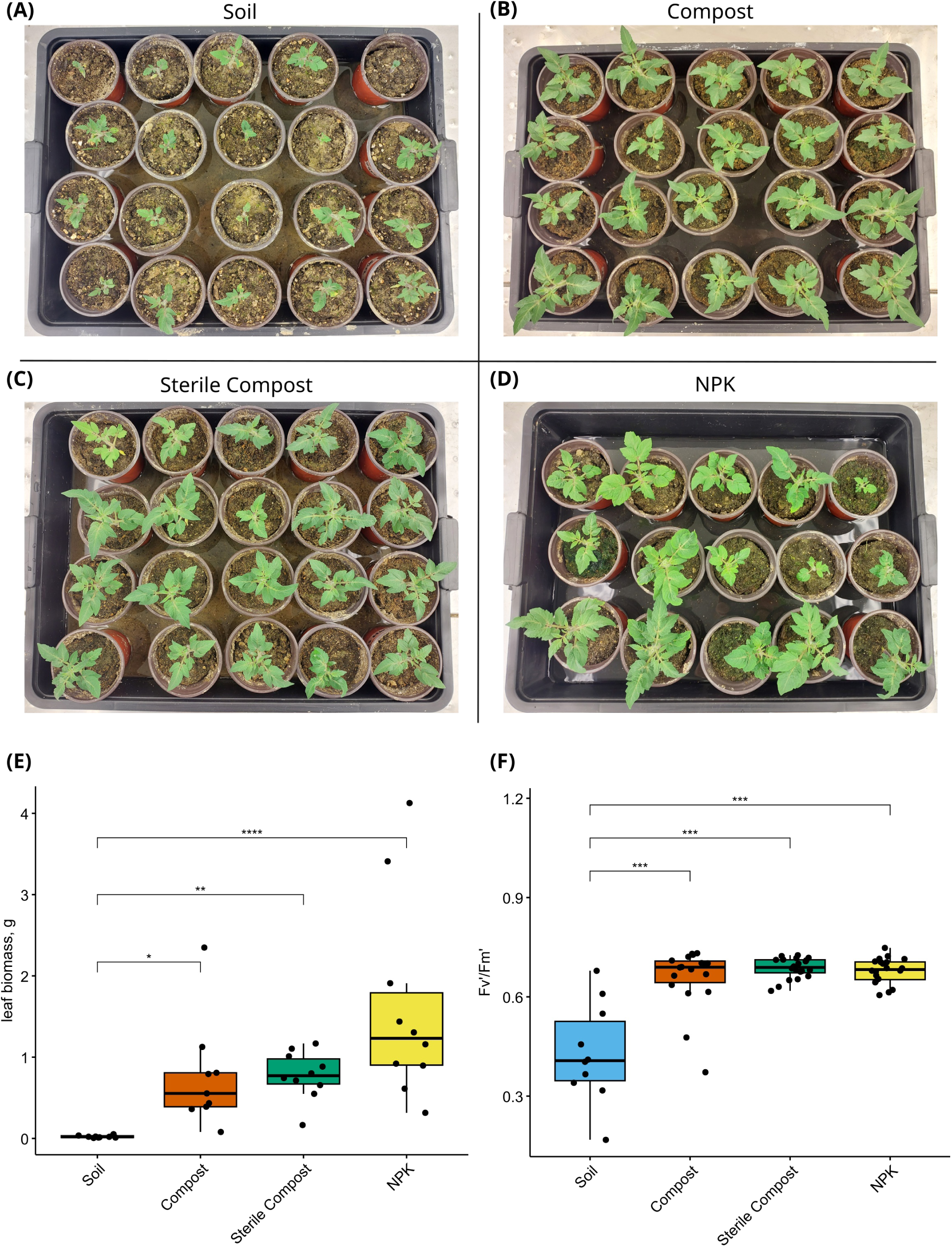
Effects of different fertilization treatments on tomato plant physiology. **(A–D)** Representative images of tomato plants after one month of growth under four soil treatments: (**A**) nutrient-poor Soil, (**B**) Compost (33 g/L), (**C**) Sterile Compost (33 g/L), and (**D**) NPK. (**E**) Boxplot showing leaf dry biomass under the four soil treatments. (**F**) Boxplot showing photosynthetic efficiency (F_v_’/F_m_’) under the different soil conditions. Statistical differences among treatments were assessed using Dunn’s test, with p-values adjusted using the Benjamini-Hochberg (BH) method. Differences were considered statistically significant at p < 0.05.

### 3.2 Compost treatment induces distinct transcriptional responses in tomato roots compared with Sterile Compost and NPK

To investigate at the molecular level the physiological differences observed among fertilization treatments, we performed RNA-sequencing analysis of tomato roots grown for one month under Compost, Sterile Compost, or chemical NPK fertilization (Figure 2). RNA-sequencing was focused on the three nutrient-rich conditions to directly compare the effects of compost-associated microbiota with those of sterilized compost and chemical fertilization.

**Figure 2.**
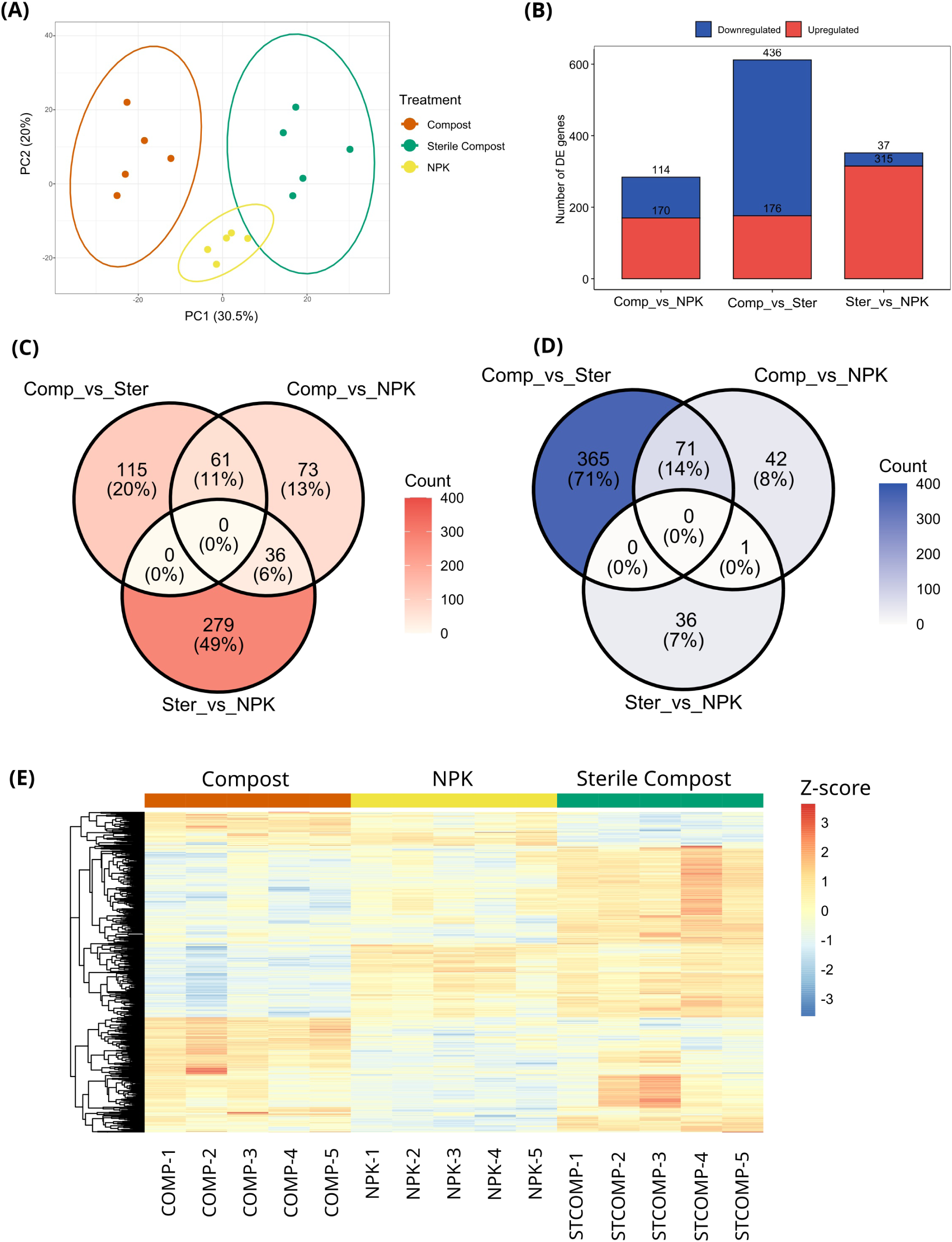
Changes in gene expression profiles of tomato roots after one month of growth under three fertilization treatments: Compost, Sterile Compost and NPK. (**A**) Principal component analysis (PCA) of the 1,000 most variable genes in tomato roots grown under the different soil treatments. Each point represents an individual biological replicate. (**B**) Stacked bar plot showing the numbers of upregulated and downregulated DEGs identified in each pairwise treatment comparison. (**C–D**) Venn diagrams illustrating the overlap of upregulated (**C**) and downregulated (**D**) genes among the pairwise comparisons of plants grown under the different soil treatments. (**E**) Heatmap of differentially expressed genes (DEGs) across the three fertilization types.

Principal Component Analysis (Fig. 2A) revealed clear separation of samples by fertilization treatment, indicating specific transcriptional responses to different nutrient inputs.

A total of 910 DEGs were identified through pairwise comparisons. There were 284, 612 and 352 DEGs in the comparisons of Compost vs NPK, Compost vs Sterile Compost, and Sterile Compost vs NPK (Fig. 2B). Especially, 170, 176, and 315 DEGs were upregulated, whereas 114, 436, and 37 were downregulated in the comparisons of Compost vs NPK, Compost vs Sterile Compost, and Sterile Compost vs NPK, respectively. A total of 73, 115 and 279 DEGs were upregulated uniquely (Fig. 2C), while the number of uniquely downregulated DEGs were 42, 365 and 36 between Compost vs NPK, Compost vs Sterile Compost, and Sterile Compost vs NPK, respectively (Fig. 2D). In addition, a hierarchical clustering analysis was performed to present a general overview of the expression pattern of DEGs, partitioning the genes into distinct functional cohorts (Fig. 2E). This approach highlighted specific transcriptomic signatures, distinguishing genes responsive solely to the chemical properties of the substrate (shared expression profiles between Compost and Sterile Compost relative to NPK) from those exclusively driven by the Compost microbiota (distinct expression changes in active Compost compared to both Sterile Compost and NPK).

To further characterize the transcriptional responses associated with Compost treatment, the expression patterns of DEGs involved in plant hormone signaling, nutrient metabolism, and stress responses in the roots were examined for each pairwise comparison among the three soil conditions (Supplementary Figure 1A-B). With regard to hormonal pathways (Fig. S1A), genes associated with gibberellin and cytokinin signaling, as well as ABA perception and metabolism, generally showed higher relative expression in Compost than in NPK and Sterile Compost. In contrast, auxin- and ethylene-related genes exhibited heterogeneous responses, with individual genes showing either higher or lower relative expression depending on the soil treatment. On the other hand, the Sterile Compost treatment was associated with higher relative expression of jasmonate-associated genes and other stress-related pathways.

Nutrient metabolism- and defense-related DEGs (Fig. S1B) likewise exhibited treatment-dependent expression patterns. Several genes involved in nitrogen and sulfur metabolism, including nitrate reductases, nitrate transporters, and glutamine synthetase, showed higher relative expression in Compost. Conversely, a subset of nutrient- and stress-associated genes, like sulfate transporters and a xyloglucan endoglucanase inhibitor displayed higher expression in Sterile Compost. NPK generally exhibited intermediate or gene-specific expression profiles.

Overall, these patterns indicate that active Compost was associated with distinct root transcriptional responses compared to both Sterile Compost and NPK. In particular, the contrast between Compost and Sterile Compost identifies candidate responses potentially linked to the compost-associated microbial community, whereas contrasts with NPK also reflect differences in substrate chemistry and nutrient availability.

To characterize the biological functions of Compost treatment-induced transcriptomes in tomato roots, GO enrichment and KEGG pathway analysis were performed to categorize upregulated and downregulated DEGs.

Of the DEGs identified in each comparison, 131 upregulated and 332 downregulated DEGs in Compost vs Sterile Compost, 129 upregulated and 99 downregulated DEGs in Compost vs NPK, and 257 upregulated and 19 downregulated DEGs in Sterile Compost vs NPK were annotated and assigned to three major GO categories (Table S3).

The dot plots in Fig. S1C-H show the significantly enriched GO terms for upregulated and downregulated genes separately within each GO ontology.

In the *Biological Processes* category (Fig. S1C-D), Compost-upregulated genes were enriched in phosphorus metabolism and responses to oxidative stress and other stimuli compared with both Sterile Compost and NPK. Carbohydrate metabolism and cell wall organization/biogenesis were uniquely enriched in Compost vs NPK, whereas nitrate-related processes, response to nitrogen compounds, and protein metabolism were specific to Compost vs Sterile Compost. Sterile Compost vs NPK was mainly enriched in photosynthesis- and electron transport-related processes. Among downregulated genes, protein and metabolic regulatory processes were enriched in Compost compared with both treatments, while photosynthesis and carbohydrate catabolism were specifically enriched in Compost vs Sterile Compost. Sterile Compost vs NPK showed only a few significant GO terms, mainly related to regulatory processes.

Regarding *Molecular Functions* (Fig. S1E-F), upregulated genes were enriched in catalytic and antioxidant activities in Compost compared with both NPK and Sterile Compost. Hydrolase and iron ion binding were specific to Compost vs NPK, whereas serine peptidase activity was specific to Compost vs Sterile Compost; electron transport and chlorophyll binding were enriched in Sterile Compost vs NPK. Among downregulated genes, most enriched terms occurred in Compost vs Sterile Compost, mainly involving cation and ATP binding and oxidoreductase activity, while Sterile Compost vs NPK showed only flavin mononucleotide (FMN) binding.

*Cellular Components* (Fig. S1G-H) contained the fewest enriched terms. Upregulated genes showed enrichment only in Sterile Compost vs NPK, mainly related to membranes and photosystems. Compost vs Sterile Compost was enriched in photosystems and thylakoids, whereas Compost vs NPK showed enrichment only for one membrane-related term.

A few pathways were significantly enriched in the KEGG analysis (Table S4). In Compost vs Sterile Compost, downregulated genes were strongly associated with photosynthesis, including antenna proteins and the Calvin cycle, as well as broader carbon and metabolic pathways, porphyrin metabolism, and the pentose phosphate pathway. Conversely, upregulated genes were enriched in nitrogen-related pathways, including nitrogen cycle and nitrogen metabolism, together with phenylpropanoid biosynthesis. In Compost vs NPK, upregulated genes were similarly enriched in nitrogen cycle and nitrogen metabolism, suggesting a consistent response associated with nitrogen-related functions in Compost. Sterile Compost vs NPK showed fewer enriched pathways, with antenna proteins and protein processing in the ER enriched among upregulated genes, while monoterpenoid biosynthesis was enriched among downregulated genes.

### 3.3 Compost supports plant physiological performance under severe water stress

The higher relative expression of genes involved in ABA signaling, nitrogen and phosphorus metabolism, and antioxidant pathways in Compost compared with Sterile Compost and the chemical fertilizer prompted us to further investigate the potential contribution of the Compost microbiota under drought stress. To this end, we performed a water stress experiment using the three soil treatments. Figure 3 shows two physiological parameters measured at the beginning of the experiment and after two weeks of water stress.

**Figure 3.**
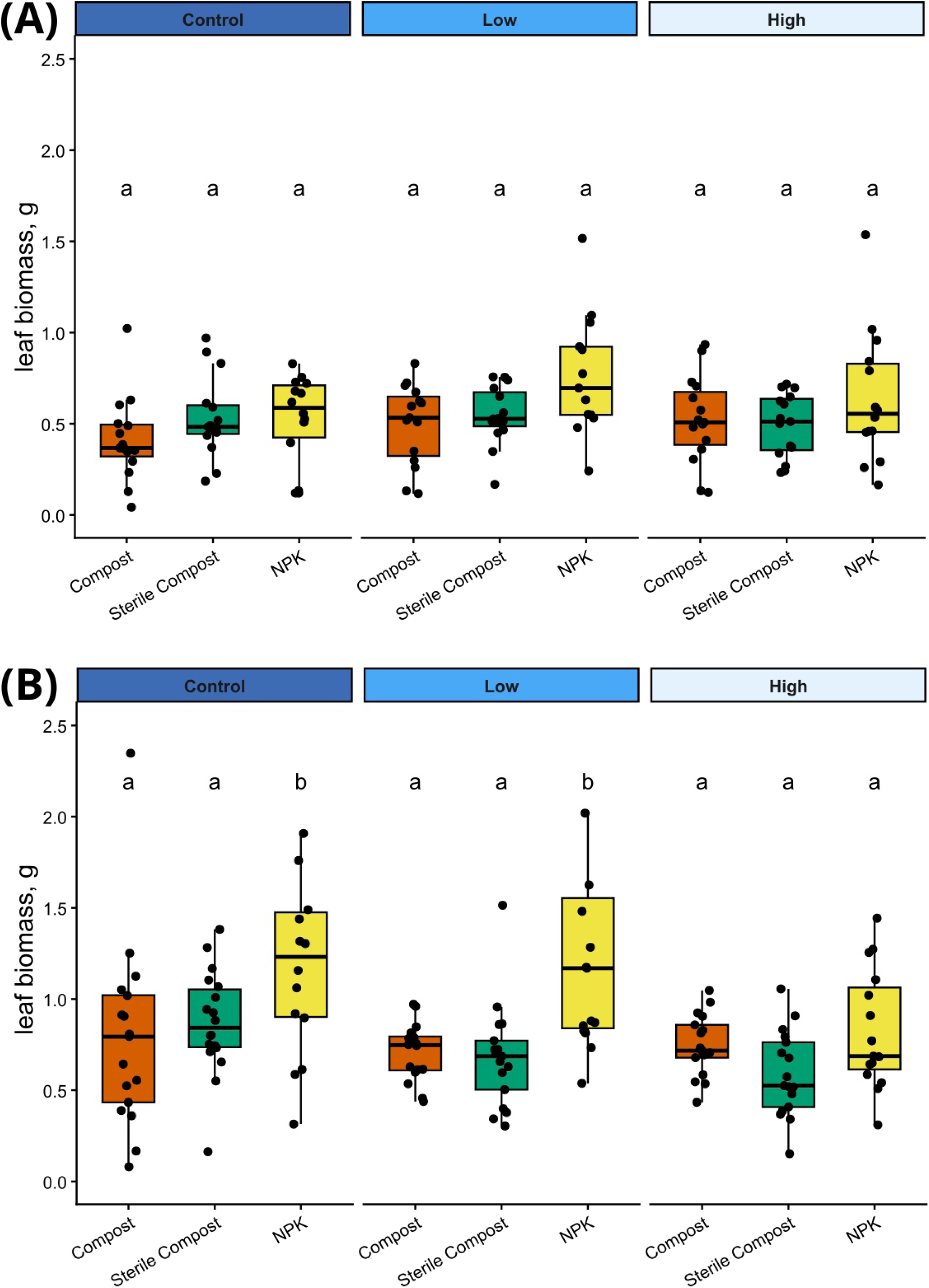
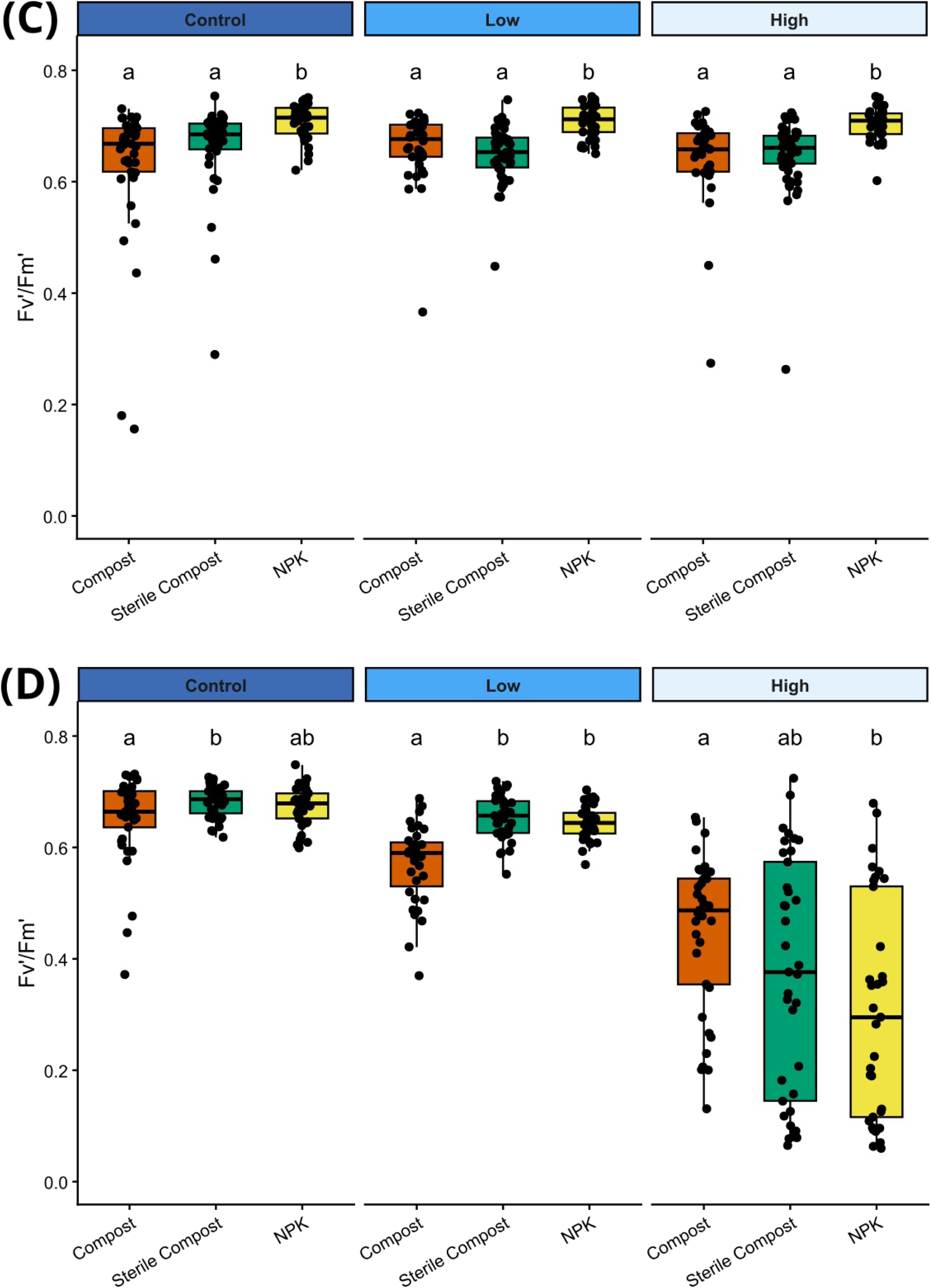
Effects of fertilization type on leaf biomass and photosynthetic efficiency of tomato plants under water stress. Physiological measurements shown according to water stress level: (**A**) Leaf dry biomass at the beginning of the water stress experiment (day 0), and (**B**) leaf biomass after 14 days of water stress. Boxplots show the distribution of individual observations; different letters indicate significant differences among fertilization types within each water-stress treatment based on BH-adjusted pairwise comparisons (α = 0.05). (**C**) Maximum quantum efficiency of photosystem II (F_v_’/F_m_’) at the beginning of the water stress experiment (day 0), and (**D**) F_v_’/F_m_’ after 14 days of water stress. Boxplots show the distribution of individual observations; different letters indicate significant differences among fertilization types within each water-stress treatment based on BH-adjusted pairwise comparisons (α = 0.05).

At day 0, leaf biomass (Fig. 3A; Fig. S2A) did not differ significantly among soil treatments. The biomass values reflected the nutritional status of the plants after three weeks of growth under the respective fertilization regimes, with Sterile Compost- and NPK-treated plants showing a tendency toward higher biomass than Compost-treated plants. Photosynthetic efficiency (Fig. 3C; Fig S2C) showed a similar trend, with NPK-treated plants exhibiting significantly higher F_v_’/F_m_’ values than plants grown in Compost or Sterile Compost.

After 14 days of water stress, a different pattern emerged. For leaf biomass (Fig. 3B), control plants showed values consistent with the initial experiment, with chemically fertilized plants exhibiting significantly higher biomass than the other treatments. Under low water stress, NPK-treated plants maintained relatively high biomass, whereas Compost- and Sterile Compost-treated plants showed a slight, but not significant, reduction. Under high water stress, no significant differences in biomass were detected among the three soil treatments. Notably, Compost was the only treatment in which biomass did not undergo a significant reduction across the water stress conditions (Fig. S2B).

Photosynthetic efficiency revealed a stronger treatment-dependent response. Within each soil treatment, F_v_’/F_m_’ decreased significantly with increasing water stress compared with the control (Fig. S2D). When comparing soil treatments within each water stress level (Fig. 3D), control plants showed comparable F_v_’/ F_m_’ values across treatments. Under low water stress, however, Compost-treated plants exhibited significantly lower photosynthetic efficiency than plants grown in Sterile Compost or NPK. Under high water stress, this pattern was reversed: Compost-treated plants retained the highest F_v_’/F_m_’ values, whereas NPK-treated plants showed the greatest reduction, with Sterile Compost plants displaying an intermediate phenotype. Overall, these results demonstrated that under severe water stress, Compost-treated plants maintained the highest photosynthetic efficiency despite not showing the highest biomass, indicating that Compost may preferentially support physiological resilience rather than growth under drought.

### 3.4 Compost amendment promotes greater water availability than Sterile Compost and NPK

To assess whether differences in substrate water availability could contribute to plant responses observed under drought, soil water retention curves and Available Water Content (AWC) were compared across the four substrate treatments (Figure 4). Unlike the drought and transcriptomic experiments, which compared nutrient-rich treatments to understand the effects of Compost microbiota relative to different nutritional inputs, the soil water retention analysis also included Soil as the substrate baseline to quantify how each amendment modified the water retention properties and AWC of the original substrate.

**Figure 4.**
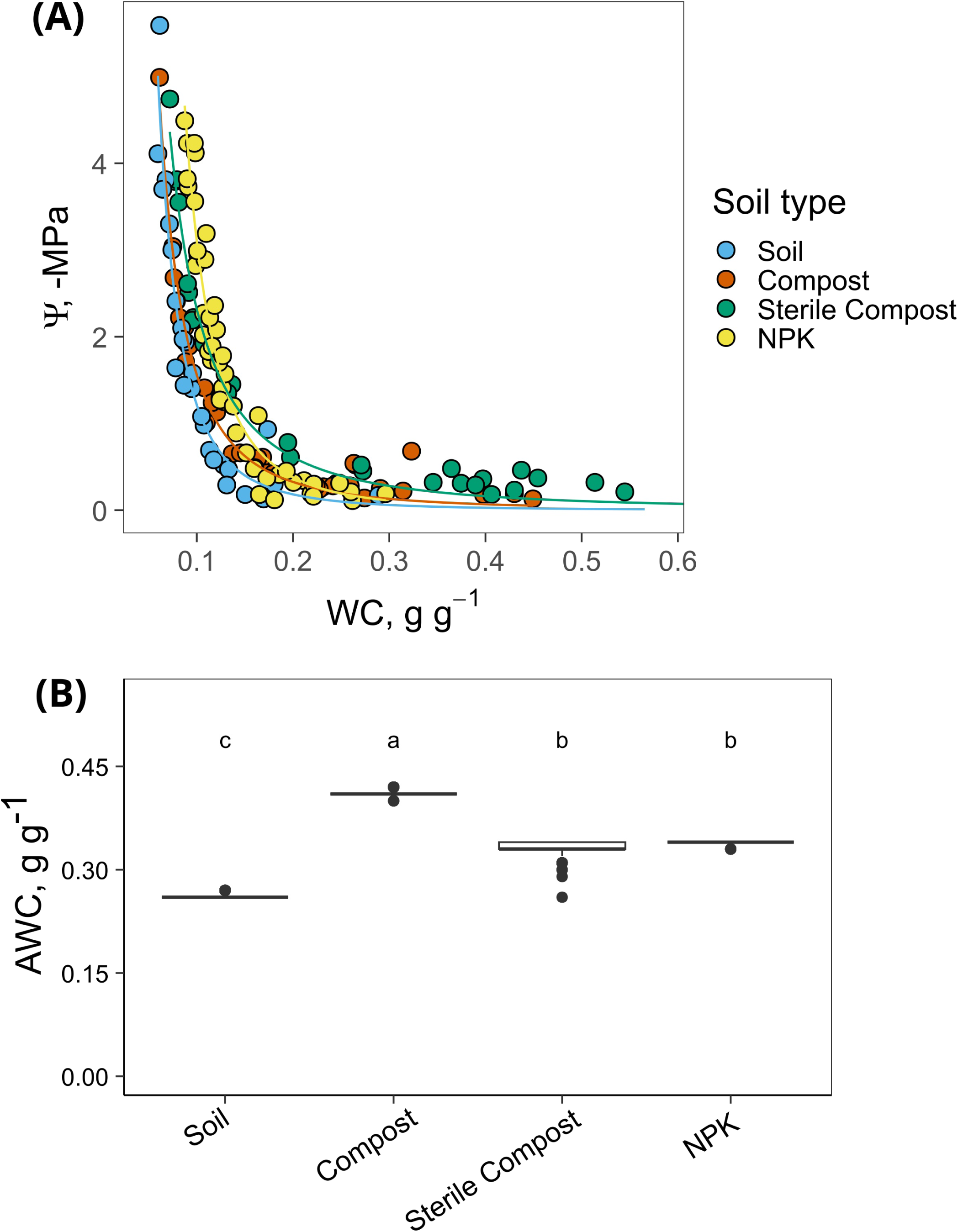
Soil water retention curves and Available Water Content (AWC) across soil treatments. (**A**) Relationship between Water Content (WC) and soil water potential (Ψ) for Soil, Compost-amended soil, Sterile Compost-amended soil, and NPK- amended soil. Points represent individual measurements, and lines represent the fitted power-curve models for each treatment. (**B**) Boxplots showing the distribution of AWC estimates obtained from 999 bootstrap resamples for each treatment. Different letters indicate differences between treatments supported by the 95% bootstrap confidence intervals of pairwise AWC differences.

The water retention curves differed among treatments, indicating distinct water retention properties of the substrates (Fig. 4A): at the same water potential, amended soils generally showed higher WC values than untreated soil, indicating a greater capacity to retain water even at lower moisture levels.

Moreover, Compost showed higher estimated AWC than both Sterile Compost and NPK, which did not differ significantly, whereas Soil showed the lowest AWC (Fig. 4B). Thus, the addition of the different amendments increased the amount of water available to plants relative to the unfertilized soil, with the greatest increase observed in Compost.

## 4. Discussion

The use of organic fertilizers such as compost in agriculture is widely recognized as a sustainable alternative to synthetic chemical inputs, as compost can improve soil physical and structural properties, enhance soil fertility, and promote the gradual release of macro- and micronutrients (Khan *et al*., 2024). These benefits extend beyond the physicochemical properties of soil, as compost application can substantially influence soil microbial diversity and functionality, which in turn contributes to crop productivity (Shu *et al*., 2022). Therefore, compost represents a promising strategy for improving crop health under environmental stress conditions. Indeed, previous studies have reported improved performance of compost-treated tomato plants under drought (González-Hernández *et al*., 2025), as well as stimulation of beneficial soil microbial communities in compost- and biochar-amended barley fields subjected to drought (Ghouili *et al*., 2026). However, the relative contribution of compost-associated microbiota and the organic matter component of compost remains poorly understood, particularly when compared with conventional chemical fertilization.

In this study, we investigated the role of compost-associated microbiota under physiological conditions, with a particular focus on root molecular responses, and under stress conditions, analyzing leaf responses.

Experimental data from the experiment with the three fertilzation types (Compost, Sterile Compost and NPK) demonstrated that NPK-treated plants displayed higher biomass than those grown with Compost or Sterile Compost, while showing comparable photosynthetic efficiency. These results showed that all three fertilization regimes were sufficient to sustain plant physiological performance, while differences in biomass among treatments suggest that the presence of the compost microbiota may not be fundamental for determining plant growth beyond nutrient supply alone. These findings are consistent with the different mechanisms of nutrient mobilization and impact on soil properties of compost and chemical fertilizers. Indeed, compost directly contributes nutrients and, at the same time, affects nutrient uptake by influencing the chemical, physical and biological properties of the soil. This combination of slow nutrient release, together with improvements in soil porosity and aggregation and the active contribution of microbial activity ensure gradual plant support, associated with a long-term productivity increase when applied to the fields (Sahoo *et al*., 2026). In contrast, conventional chemical fertilizers provide nutrients more rapidly, as their highly water-soluble mineral salts enable a quick increase in plant growth and yield, but their repeated application may negatively affect soil health by altering soil organic carbon and microbial communities over time (Howe *et al*., 2024).

At the molecular level, the RNA-sequencing analysis revealed a different pattern from that observed at the phenotypic level.

Although Compost, Sterile Compost, and NPK provided comparable macronutrient inputs, PCA and differential expression analyses revealed substantial differences in gene expression across treatments. This finding suggests that the plant response to compost cannot be explained solely by its nutrient content and that its associated microbial community also contributes to shaping root molecular responses.

Compost treatment upregulated several genes involved in hormonal pathways, as well as nutrient uptake and antioxidant responses compared to Sterile Compost and NPK. The transcriptional response revealed a particularly strong association with nitrogen metabolism: genes involved in nitrate reduction, nitrate transport and nitrogen assimilation, including nitrate reductases, nitrate transporters and glutamine synthetase, showed higher expression in Compost than in the other treatments. This provides evidence for a contribution of the living component of compost. Sterilization removes the viable microbial community while retaining the organic and nutritional components of the compost, so the Compost–Sterile Compost comparison suggests that the microbial component of compost may contribute to the regulation of nitrogen acquisition or assimilation, whereas the Compost–NPK comparison indicates that this response is not simply explained by the macronutrient supply. These results provide molecular evidence that the microbiota associated with urban biowaste compost contributes to plant health and physiological performance, extending previous agronomic findings obtained using, for instance, onion-residue and cattle-manure compost in field-grown tomato (Pellejero *et al*., 2021) and sewage-sludge and green-waste composts in maize (Zapałowska and Jarecki, 2024). Indeed, our work distinguishes microbial effects from the physico-chemical effects of urban biowaste compost by comparing sterile and non-sterile compost.

In the drought stress experiment, leaf biomass and photosynthetic efficiency, used as proxies for tomato plant performance, showed treatment-dependent responses that varied with the severity of water stress. Under low water stress, plants grown with Sterile Compost and NPK maintained higher physiological performance than those in the Compost treatment, which instead was able to maintain higher physiological parameters under high water stress. Compost is known to harbor beneficial microorganisms (Tondello *et al*., 2022; Aguilar-Paredes *et al*., 2023), and its application can shape rhizosphere microbial communities (De Brito Alvarez *et al*., 1995; Wang *et al*., 2022). More broadly, beneficial plant-associated microorganisms can enhance plant growth and improve tolerance to a range of biotic and abiotic stresses (De Andrade *et al*., 2023), suggesting that Compost-associated microbiota may have contributed to the improved plant response under severe drought. This is also consistent with the transcriptomic data, which showed higher expression of genes involved in ABA perception and metabolism in Compost-treated plants, potentially indicating enhanced regulation of a central pathway controlling plant responses to water limitation. However, further experiments examining root molecular responses directly under drought will be required to establish whether these changes contribute to the enhanced drought resilience observed in Compost-treated plants.

Compost showed greater water retention and the highest AWC, indicating that its addition increased the amount of water potentially available to plants. This is consistent with previous studies showing that compost improves soil structure, aggregation, and pore-size distribution, thereby enhancing water retention (Bondì *et al*., 2022). Indeed, the higher organic matter content associated with compost can increase water storage and reduce water losses through drainage and evaporation (Iraji *et al*., 2025). These improvements in physico-chemical properties are further enhanced by microbial activity, as microorganisms can contribute to soil structural stability and reduced compaction by secreting sticky organic compounds or through the development of fungal hyphal networks (Rillig and Mummey, 2006). In contrast, the lower water-holding capacity observed in the other amendments may reflect treatment-specific features. In Sterile Compost, the sterilization process may have altered the physical properties of the substrate while drastically reducing its biological component, potentially limiting microbially mediated aggregation and structural stabilization (Berns *et al*., 2008). In NPK, the lower water capacity may instead be related to changes in bulk density and porosity associated with the absence of organic matter and the increased presence of salts (Acar *et al*., 2025). Overall, the synergy between the greater AWC provided by Compost and the contribution of its microbial community to soil structural stability may have enhanced water availability under drought, thereby contributing to the better maintenance of physiological performance observed under severe drought conditions.

In conclusion, our results demonstrate that compost from biowaste can influence plant performance beyond its nutritional contribution, with its associated microbiota playing an important role in shaping root molecular responses and plant adaptation to drought stress. Including sterilized compost and an NPK treatment supplying macronutrients at levels equivalent to those provided by compost allowed us to partially disentangle the contribution of the living microbial component from the nutritional and physico-chemical effects of compost. The synergy between compost-associated microbial activity and improved substrate water-holding capacity may be particularly relevant under severe drought, helping to maintain plant physiological performance. These findings highlight the potential of compost as a sustainable soil amendment and support further investigation of the specific microbial and molecular mechanisms underlying its effects on plant resilience.

## Authors’ contributions

Conceptualization, SN and RC; methodology, RC, FP and SN; formal analysis, RC, FP and SRS; investigation, GG, RC and LG; resources, SN, BB, SRS, TB and WZ; data curation, RC and SRS; supervision, SN and BB; funding acquisition, SN and BB; writing - original draft preparation, RC; writing - review and editing, SN, RC, BB and FP. All authors reviewed and approved the final version of the manuscript.

## Acknowledgements

We thank Rebecca Montanari, MSc, for her contribution to the assessment of soil available water content; Annalaura Masiero and Sebastiano Fossaluzza, BSc, for their assistance with sampling and physiological measurements; and Prof. Marco Giovannetti (University of Turin, Italy) and Prof. Dr. Gabriele Berg (Graz University of Technology, Graz, Austria), doctoral advisors of RC, for their valuable suggestions.

## Data availability statement

The sequence dataset generated during the current study is available in the NCBI SRA database under the BioProject PRJNA1545275.

The supplementary tables are available on Zenodo (https://doi.org/10.5281/zenodo.23403897).

