## Supplementary Figures for "Compost-associated microbiota modulates root transcriptional responses and enhances tomato drought resilience"

### Supplementary Tables

The supplementary tables are available on Zenodo (*https://doi.org/10.5281/zenodo.23403897*).

**Table S1.** Soil physico-chemical properties

**Table S2.** Compost physico-chemical properties

**Table S3.** Summary of Gene Ontology annotations for up- and down-regulated genes identified in pairwise substrate comparisons.

**Table S4.** KEGG pathway enrichment analysis of up- and down-regulated genes across pairwise substrate comparisons.

### Supplementary Figures

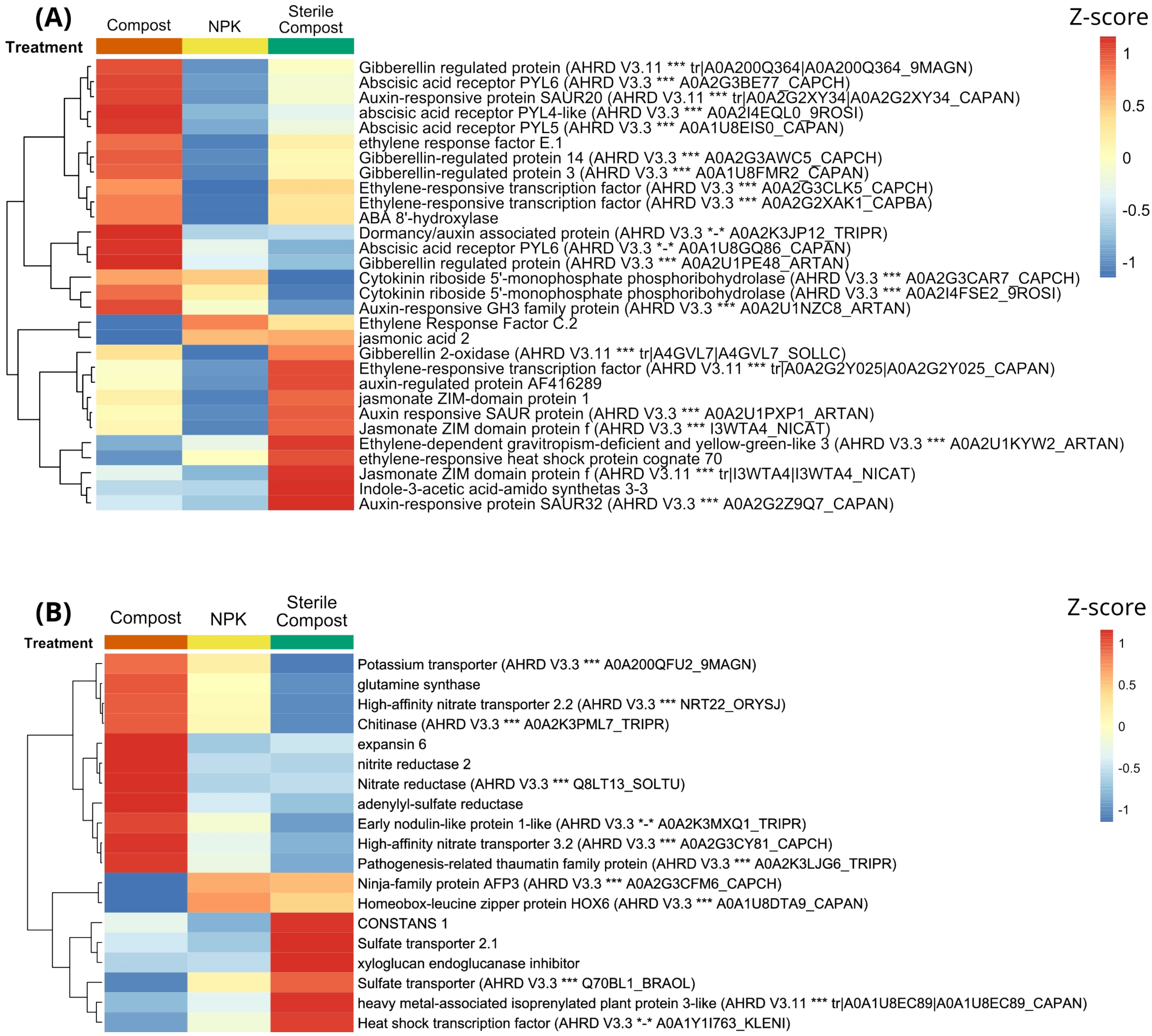


**Figure S1. Transcriptional responses associated with hormones, nutrient metabolism, defense, and Gene Ontology enrichment in tomato roots in different soil conditions. (A)** Heatmap of differentially expressed genes (DEGs) related to hormone signaling across the three soil conditions: Compost, Sterile Compost, and NPK. **(B)** Heatmap of DEGs associated with nutrient metabolism and defense responses across the same soil conditions. For both heatmaps, expression values are presented as Z-scores, calculated by centering and scaling the normalized expression values for each gene across samples. **(C–D)** Comparative dot plots of enriched GO Biological Processes across the three pairwise comparisons, showing upregulated genes (**C**) and downregulated genes (**D**). **(E–F)** Comparative dot plots of enriched GO Molecular Functions, showing upregulated genes (**E**) and downregulated genes (**F**). **(G–H)** Comparative dot plots of enriched GO Cellular Components, showing upregulated genes (**G**) and downregulated genes (**H**). For all comparative dot plots, enrichment is shown across the three pairwise comparisons: Compost vs NPK, Compost vs Sterile Compost, and Sterile Compost vs NPK. Dot size represents the Gene Ratio, calculated as the number of significant genes associated with a given GO term divided by the total number of unique significant genes included in the corresponding enrichment analysis. Dot colour indicates the statistical significance of GO enrichment, with lower P values indicating stronger enrichment.

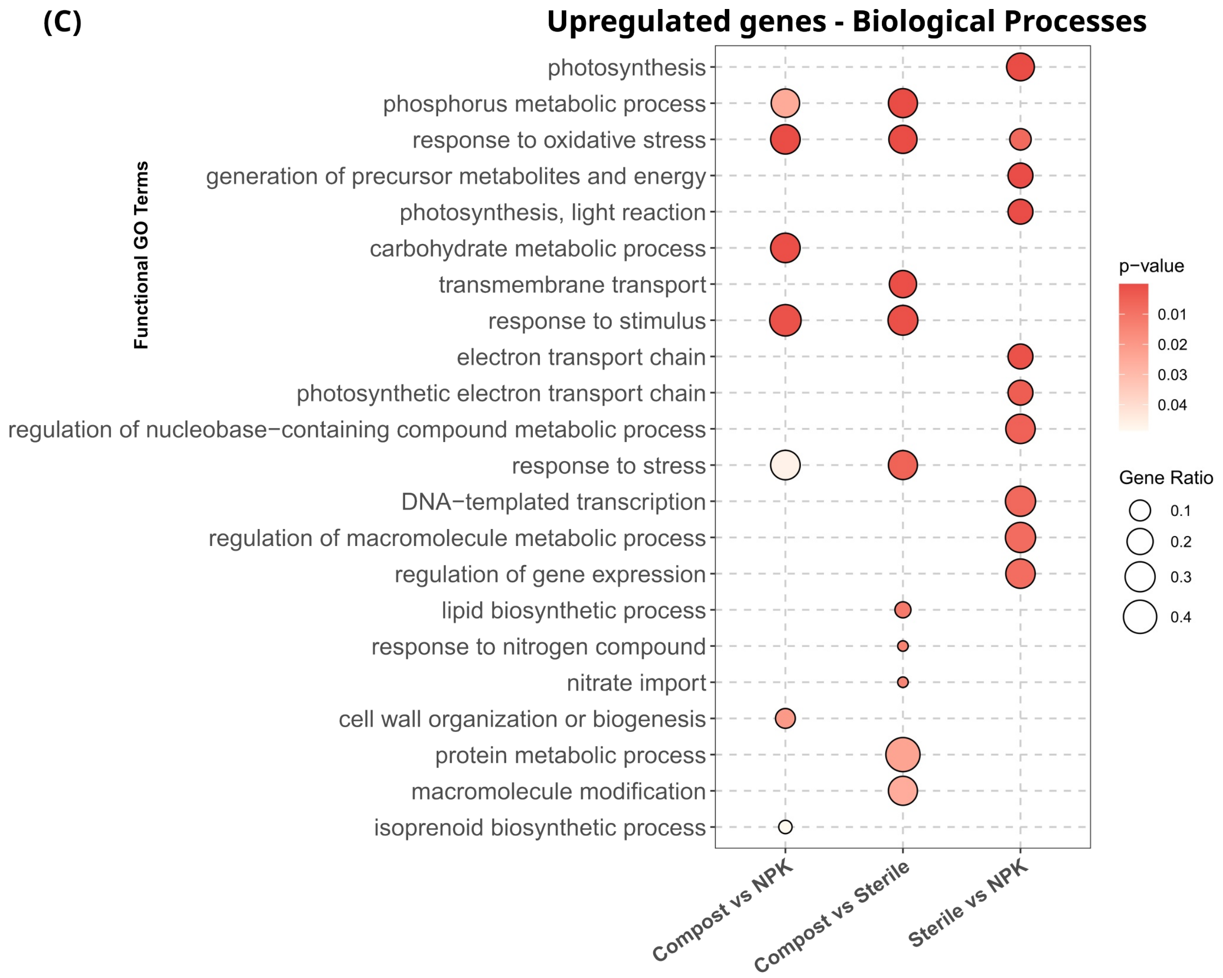
(C)
Upregulated genes - Biological Processes
Functional GO Terms
photosynthesis
phosphorus metabolic process
response to oxidative stress
generation of precursor metabolites and energy
photosynthesis, light reaction
carbohydrate metabolic process
transmembrane transport
response to stimulus
electron transport chain
photosynthetic electron transport chain
regulation of nucleobase−containing compound metabolic process
response to stress
DNA−templated transcription
regulation of macromolecule metabolic process
regulation of gene expression
lipid biosynthetic process
response to nitrogen compound
nitrate import
cell wall organization or biogenesis
protein metabolic process
macromolecule modification
isoprenoid biosynthetic process
Compost vs NPK
Compost vs Sterile
Sterile vs NPK
p−value
0.01
0.02
0.03
0.04
Gene Ratio
0.1
0.2
0.3
0.4

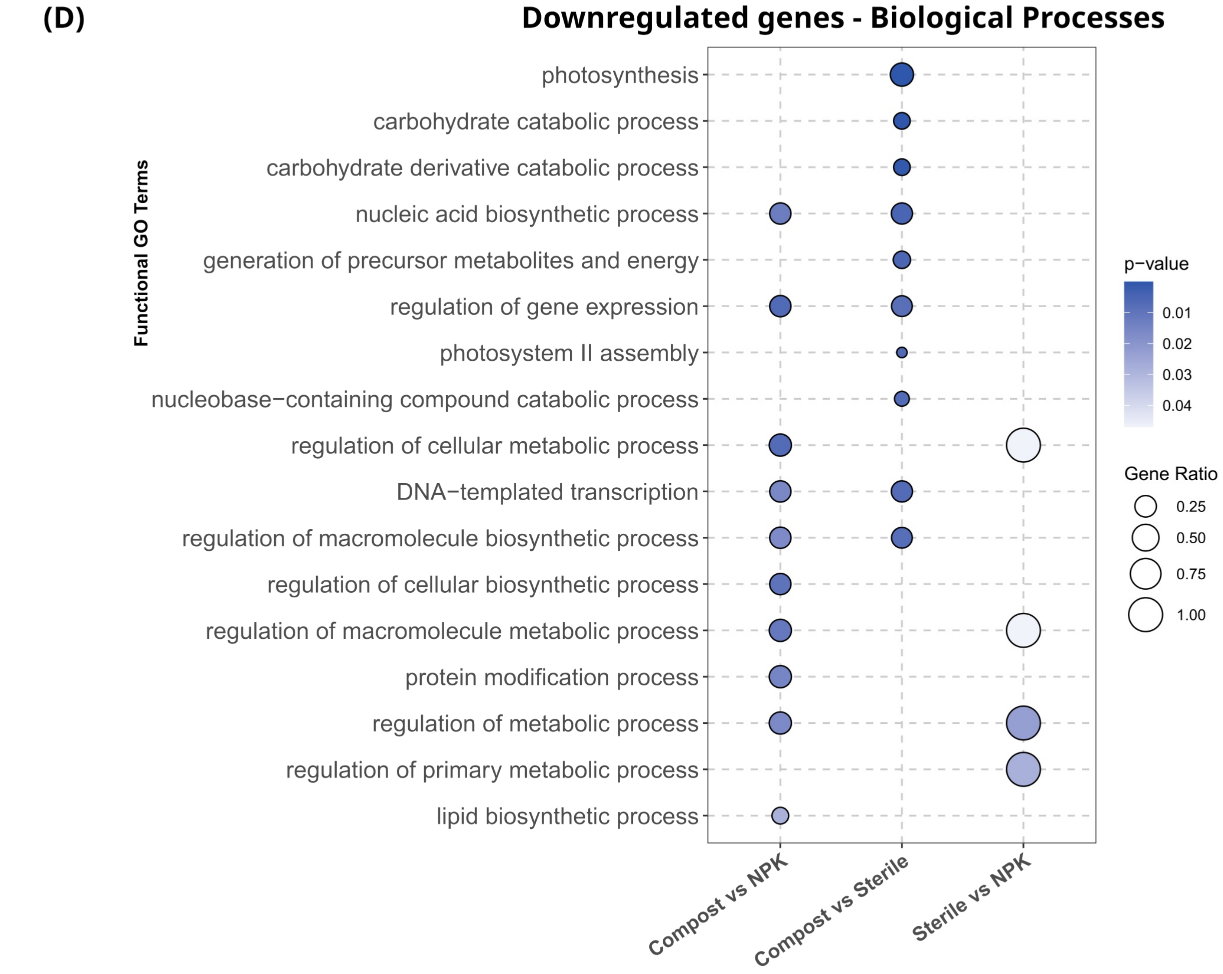
(D)
Downregulated genes - Biological Processes
Functional GO Terms
photosynthesis
carbohydrate catabolic process
carbohydrate derivative catabolic process
nucleic acid biosynthetic process
generation of precursor metabolites and energy
regulation of gene expression
photosystem II assembly
nucleobase−containing compound catabolic process
regulation of cellular metabolic process
DNA−templated transcription
regulation of macromolecule biosynthetic process
regulation of cellular biosynthetic process
regulation of macromolecule metabolic process
protein modification process
regulation of metabolic process
regulation of primary metabolic process
lipid biosynthetic process
Compost vs NPK
Compost vs Sterile
Sterile vs NPK
p−value
0.01
0.02
0.03
0.04
Gene Ratio
0.25
0.50
0.75
1.00

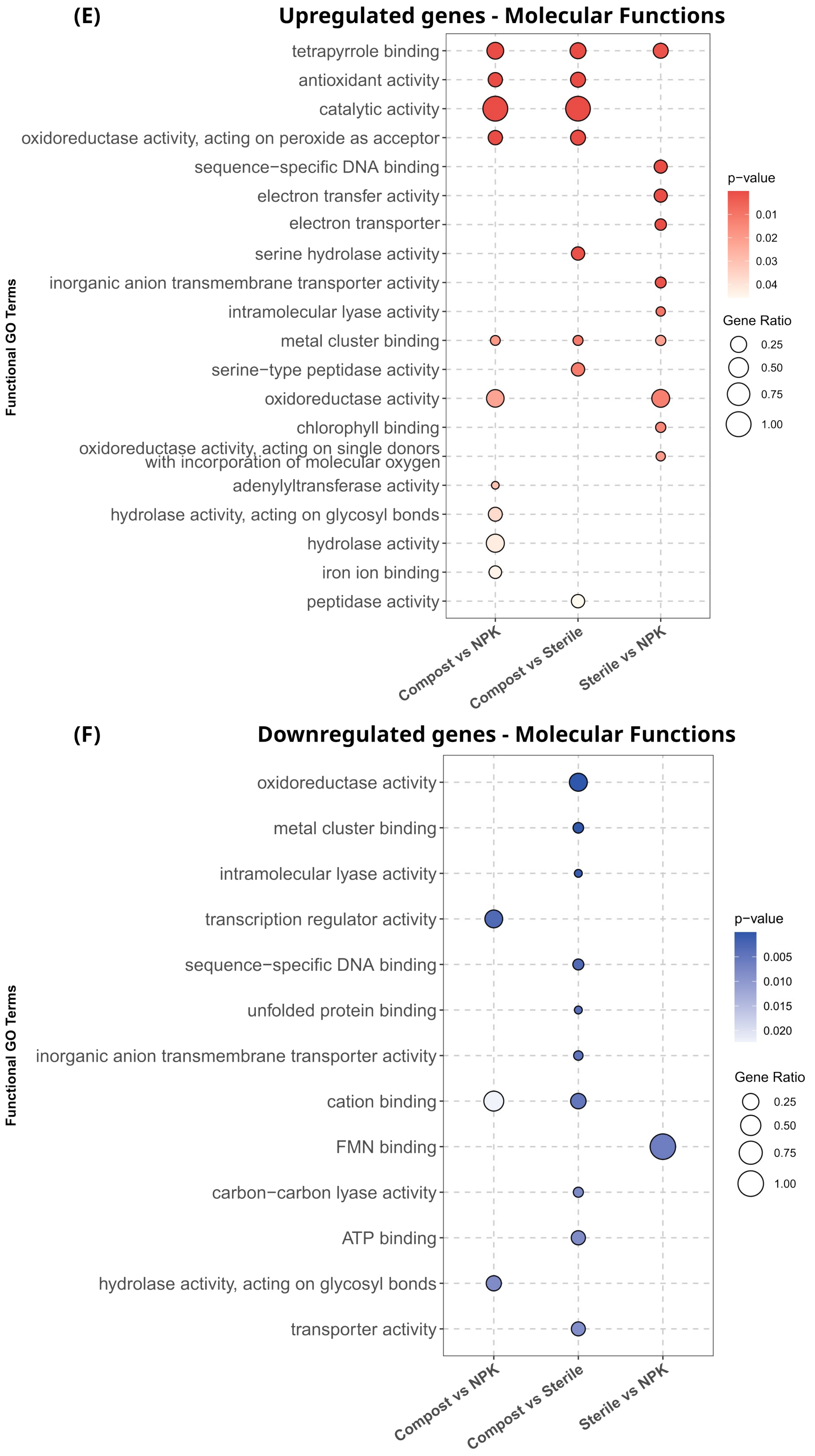

(E)
Upregulated genes - Molecular Functions
tetrapyrrole binding
antioxidant activity
catalytic activity
oxidoreductase activity, acting on peroxide as acceptor
sequence−specific DNA binding
electron transfer activity
electron transporter
serine hydrolase activity
inorganic anion transmembrane transporter activity
intramolecular lyase activity
metal cluster binding
serine−type peptidase activity
oxidoreductase activity
chlorophyll binding
oxidoreductase activity, acting on single donors with incorporation of molecular oxygen
adenylyltransferase activity
hydrolase activity, acting on glycosyl bonds
hydrolase activity
iron ion binding
peptidase activity
Functional GO Terms
Compost vs NPK
Compost vs Sterile
Sterile vs NPK
p−value
0.01
0.02
0.03
0.04
Gene Ratio
0.25
0.50
0.75
1.00
(F)
Downregulated genes - Molecular Functions
oxidoreductase activity
metal cluster binding
intramolecular lyase activity
transcription regulator activity
sequence−specific DNA binding
unfolded protein binding
inorganic anion transmembrane transporter activity
cation binding
FMN binding
carbon−carbon lyase activity
ATP binding
hydrolase activity, acting on glycosyl bonds
transporter activity
Functional GO Terms
Compost vs NPK
Compost vs Sterile
Sterile vs NPK
p−value
0.005
0.010
0.015
0.020
Gene Ratio
0.25
0.50
0.75
1.00

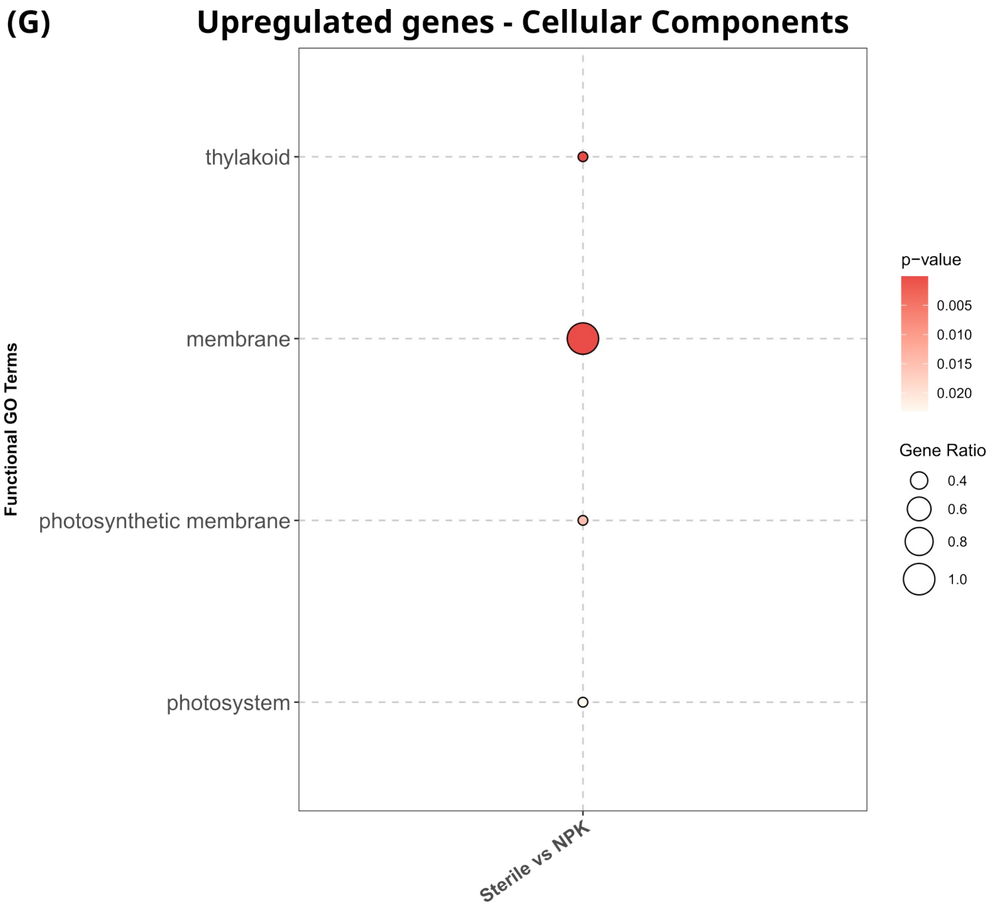
(G)
Upregulated genes - Cellular Components
Functional GO Terms
thylakoid
membrane
photosynthetic membrane
photosystem
Sterile vs NPK
p−value
0.005
0.010
0.015
0.020
Gene Ratio
0.4
0.6
0.8
1.0

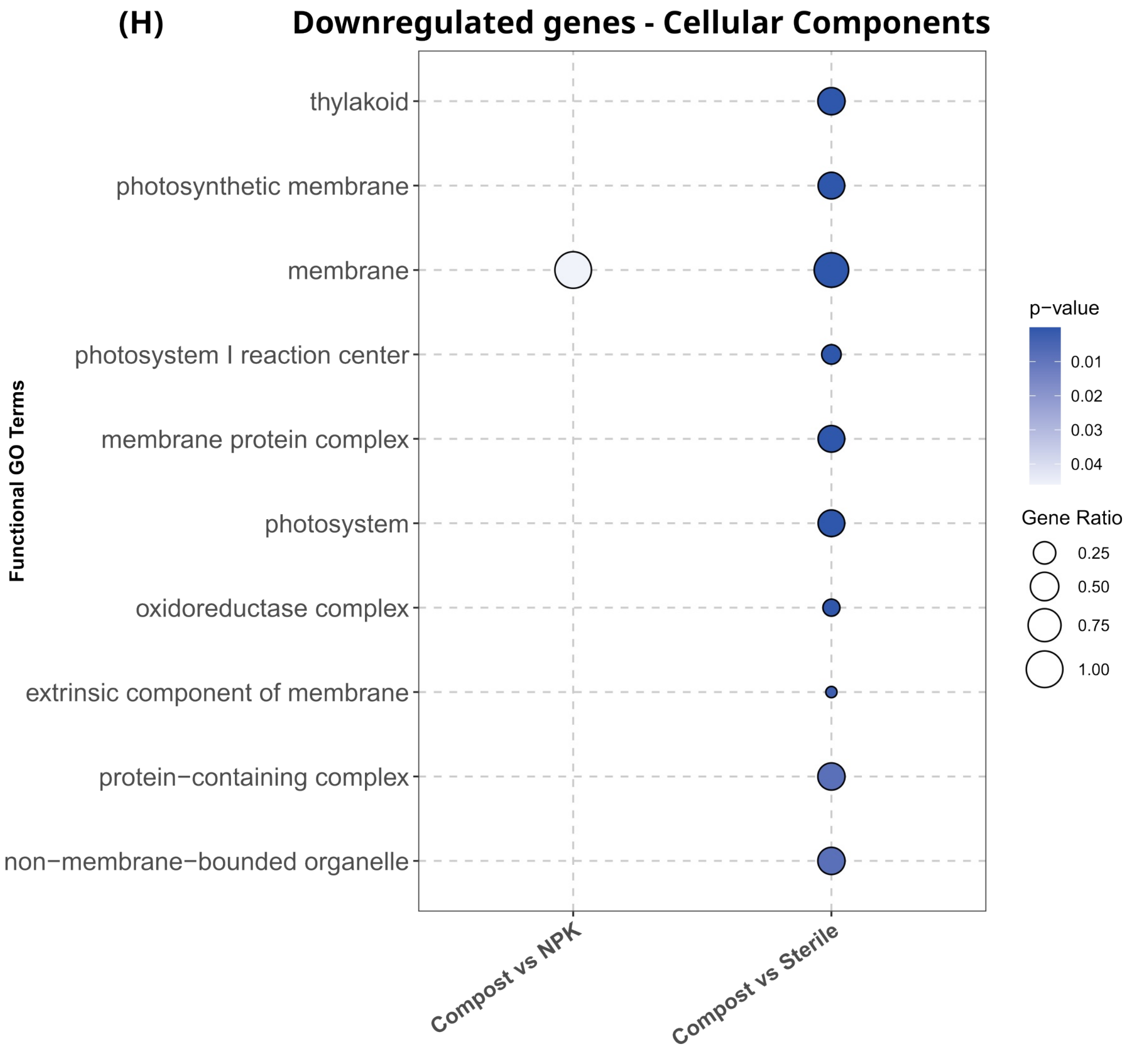
(H)
Downregulated genes - Cellular Components
Functional GO Terms
thylakoid
photosynthetic membrane
membrane
photosystem I reaction center
membrane protein complex
photosystem
oxidoreductase complex
extrinsic component of membrane
protein−containing complex
non−membrane−bounded organelle
Compost vs NPK
Compost vs Sterile
p−value
0.01
0.02
0.03
0.04
Gene Ratio
0.25
0.50
0.75
1.00

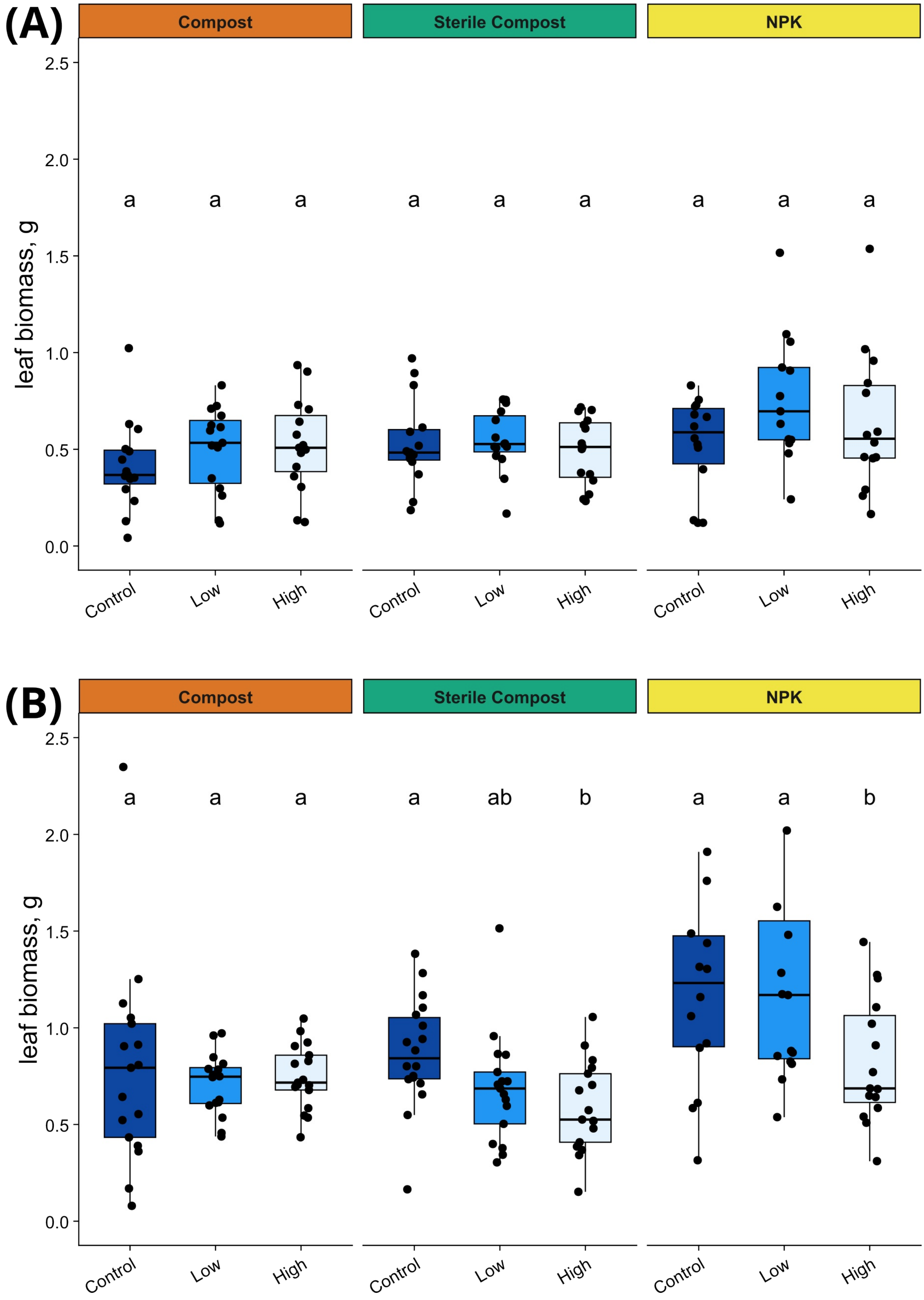


**Figure S2. Effects of water stress on biomass and photosynthetic efficiency of tomato plants grown under different fertilization conditions.** Physiological measurements shown according to soil type: **(A)** Leaf dry biomass at the beginning of the water stress experiment (day 0), and **(B)** leaf biomass after 14 days of water stress. Boxplots show the distribution of individual observations; different letters indicate significant differences among fertilization types within each water-stress treatment based on BH-adjusted pairwise comparisons (α = 0.05).

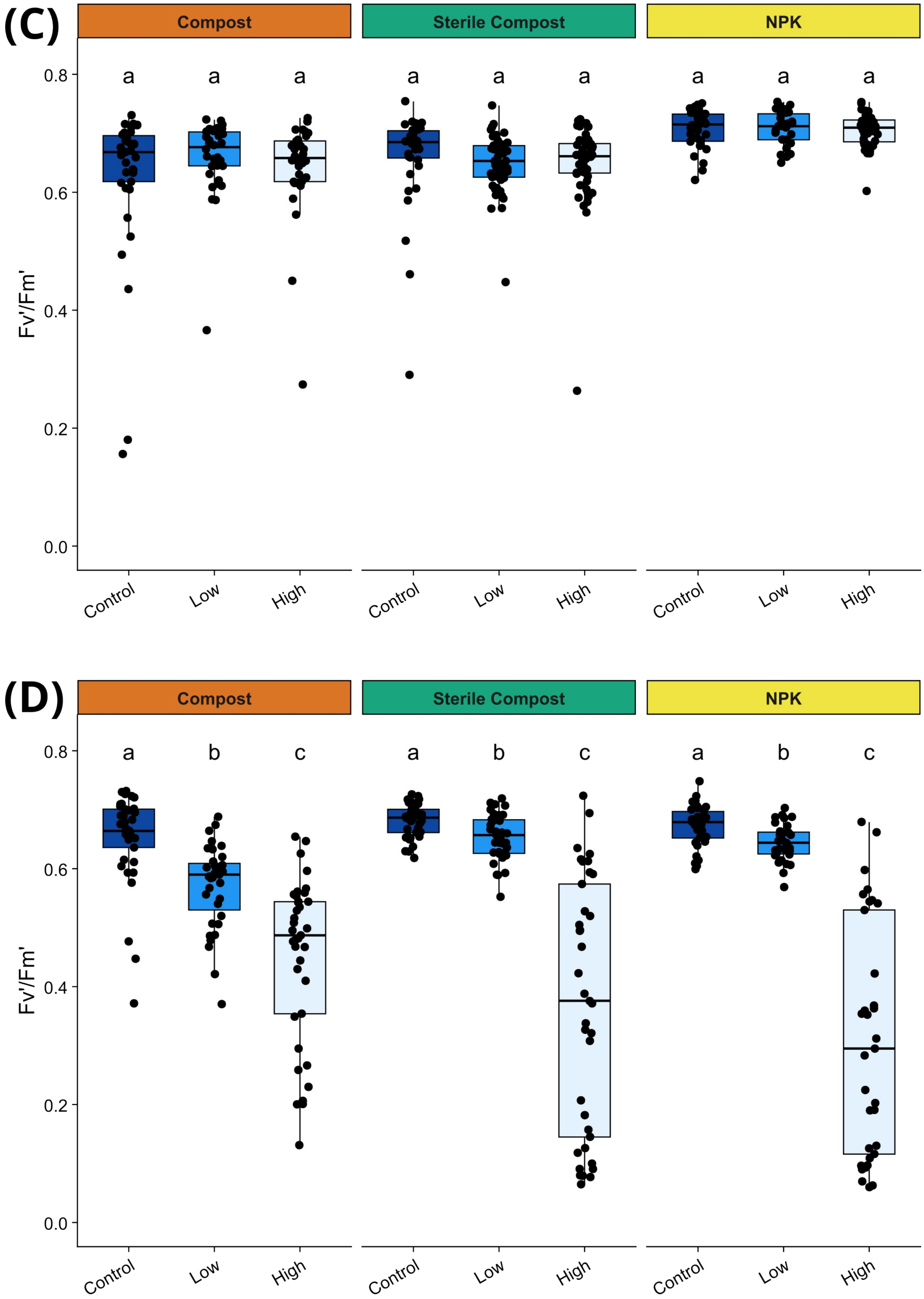


**Figure S2. Effects of water stress on biomass and photosynthetic efficiency of tomato plants grown under different fertilization conditions.** Physiological measurements shown according to soil type: **(C)** Maximum quantum efficiency of photosystem II ($F_v'/F_m'$) at the beginning of the water stress experiment (day 0), and **(D)** $F_v'/F_m'$ after 14 days of water stress. Boxplots show the distribution of individual observations; different letters indicate significant differences among fertilization types within each water-stress treatment based on BH-adjusted pairwise comparisons ($\alpha = 0.05$).
